# Everyone Can Meditate: Characterizing a Personalized Connectomic State Space among Meditation Groups and Non-meditators

**DOI:** 10.1101/2020.06.19.162461

**Authors:** Jacob van Doorn, Mengqi Xing, B. Rael Cahn, Arnaud Delorme, Olusola Ajilore, Alex D. Leow

## Abstract

Alterations in brain connectivity has been shown for many disease states and groups of people from different levels of cognitive training. To study dynamic functional connectivity, we propose a method for a personalized connectomic state space called Thought Chart. Experienced meditators are an interesting group of healthy subjects for brain connectivity analyses due to their demonstrated differences in resting state dynamics, and altered brain connectivity has been implicated as a potential factor in several psychiatric disorders. Three distinct techniques of meditation are explored: Isha Yoga, Himalayan Yoga, and Vipassana, as well as a meditation-naïve group of individuals. All individuals participated in a breath awareness task, an autobiographical thinking task, and one of three different meditation practices according to their expertise, while being recorded by a 64-electrode electroencephalogram (EEG). The functional brain connectivity was estimated using weighted phase lag index (WPLI) and the connectivity dynamics were investigated using a within-individual formulation of Thought Chart, a previously proposed dimensionality reduction method which utilizes manifold learning to map out a state space of functional connectivity. Results showed that the two meditation tasks (breath awareness task and own form of meditation) in all groups were found to have consistently different functional connectivity patterns relative to those of the instructed mind-wandering (IMW) tasks in each individual, as measured using the Hausdorff distance in the state space. The specific meditation state was found to be most similar to the breath awareness state in all groups, as expected in these meditation traditions which all incorporate breath awareness training in their practice trajectory. The difference in connectivity was found to not be solely driven by specific frequency bands. These results demonstrate that the within-individual form of Thought Chart consistently and reliably separates similar tasks among healthy meditators and non-meditators during resting state-like EEG recordings. Unexpectedly, we found the dissimilarity between breath awareness/meditation and IMW, measured via Hausdorff distance, regardless of meditation experience or tradition, with no significant group differences.

## 1. Introduction

The ability to construct and analyze the state space of mental activity, both in resting state and task states, could be a useful tool in the future of brain connectivity research. Altered patterns in functional brain connectivity have been demonstrated in many mental health disorders including major depressive disorder [4], bipolar depression [25, 33], schizophrenia [16], social anxiety disorder [29, 12], and may be a biomarker for those and other psychiatric diseases [48]. In addition to evidence that disease states may be associated with altered brain connectivity patterns, there is extensive evidence that brain connectivity is altered in association with a number of beneficial training regimens such as those practiced by experienced meditators [6, 20, 26]. Recently, our group has investigated dynamic EEG connectivity to construct such a metric space of the underlying temporal connectivity yielding a state space of the mind called the Thought Chart [62]. After constructing such a state space, Xing et al. were able to identify group differences in social anxiety using the temporal pattern of navigation in the Thought Chart [63]. As we have shown the Thought Chart to be effective in comparing patients and healthy groups, we new demonstrate its utility to explore specific mental states in healthy subjects.

The brain’s connectivity can be analyzed on a global or local scale by applying concepts from graph theory [13]. Brain networks, both structural and functional, have been recently shown to have specific graph theory metrics, including small-worldness, high connectivity hubs, and modularity [8]. There are also well documented brain networks whose intra-network connectivities have been shown to be altered in various disease states such as the Default Mode Network (DMN) [7]. DMN intra-network connectivity has been negatively correlated with frontal theta EEG activity in the resting state [43] and is reflected in its structural connectivity as well [17]. Connectomics, as a graph-theoretical tool of neuroscience that analyzes the structure and function of either static or dynamic brain connectivity, aims to shed light on the interplay of many such networks [49, 48]. The use of electroencephalography (EEG) to build connectomes has been utilized for over a decade [44] and continues to provide insights into cognition and psychiatric disorders [22]. Dynamic functional connectivity (dFC), which involves how a connectome may change over small time scales, is a promising and more complex alternative to static connectivity [21].

The state space that this paper explores are those associated with meditation, which has consistently been found to have unique functional properties, as measured by EEG [55, 9, 3], MEG [36], fMRI [52, 23, 20, 5], DTI [31], and PET [30]. Meditation has also been found to be a possible treatment option for several mental disorders [46], for example it demonstrates efficacy in the treatment of recurrent depression, binge eating disorder, PTSD, and chronic pain [60, 38, 65]. Experienced meditators have also been shown to have differing resting state dynamics compared to meditation-naïve individuals, which make them an interesting group to include in resting state dynamic functional connectivity studies [5, 6]. While meditation has been previously studied using neuroimaging methods, there is still a gap in the literature on the differences between meditation traditions, and to what extent those differences can be observed [15]. Meditation techniques are often reduced to a spectrum on which they can be classified, from focused attention to open monitoring [32]. Broadly, focused attention meditation involves concentration on an object, such as one’s breath, a mantra, or a visualization; whereas open monitoring meditation practice involves staying attentive moment by moment to anything that occurs in experience without focusing on any explicit object [32]. Most traditions of meditation incorporate both of these styles to some degree in their practice trajectories, although some will focus more on one form or the other. Three schools of meditation were included in order to have a representative sample of the varying types of meditation found in different traditions. A more detailed description of the meditation techniques used in this study is provided section 2.2. Each individual was given two different general tasks to perform while EEG was recorded: meditation tasks vs. instructed mind-wandering.

The aim of this study is to use the EEG recordings from these three groups of experienced meditators and meditation-naïve individuals to characterize and validate a state space of the connectomes produced during meditative tasks and instructed mind-wandering, called an individualized Thought Chart. The state space contains the information of transitions from one state to another, whether that be a direct transition or indirect through intermediate states, and enables analysis of the dynamics of a system via the graph theory tools mentioned above. Thought Chart is a tool that is well-suited to exploring subtle differences in the dynamic changes in functional connectivity of different traditions of meditation. By using only healthy subjects, with variation in resting state characteristics due to their meditation experience [52, 23], we seek to establish the feasibility of the individualized Thought Chart as a personalized method for mapping the functional state space of mental activity in healthy individuals. For all groups in this study, it was hypothesized that the two meditative tasks (breath awareness and other specific meditations, as specified below) would occupy a separate area of the state space from the instructed mind-wandering task, while tasks of the same type would occupy the same area of the state space.

## 2. Materials and Methods

### 2.1. Data collection and preprocessing

Thirty-two healthy control subjects and 20 meditators from the Vipassana tradition, 27 meditators from Himalayan Yoga tradition, and 20 meditators from Isha Yoga tradition were recruited and data collection took place at the Meditation Research Institute in Rishikesh, India. Subjects were compensated and reimbursed for travel, accommodation, and meals. The project was approved by the local Indian ethical committee and the IRB of the University of California San Diego (IRB #090731). Control subjects were selected for participation in this study based on their age, gender, and lack of meditation experience. For the “meditation condition,” they were instructed to focus on the sensation of their breath for the duration of the recording period (“keep paying attention to the sensations of the breath, both of the inhalation and the exhalation. If your mind starts to wander, please bring it back to your breath.”)

The subjects were recorded using a 64 + 8 channel Biosemi Active-Two amplifier system with a 10–20 headcap standard 64-channel cap also from Biosemi. External electrodes recorded the mastoids and electrooculogram (EOG). During the tasks, subjects sat on a floor blanket or a chair, depending on their preference, and performed each task (meditation vs. instructed mind-wandering) for 20 minutes. For the meditation block, participants engaged first in ten minutes of focused attention on the breath and then at the 10 minute mark a bell sound was presented via speakers in the experimental room and participants transitioned to practicing the specific mediation practice with which they were most expert. The meditation-naïve participants were told to continue the meditation on their breath after the bell, as they had no history of expertise with any other specific meditative practice. For the instructed mind-wandering task participants were asked to mentally review a series of emotionally neutral memories of their own choosing. After the first ten minutes of the mind-wandering recording a bell was played as well, for consistency with the meditation period, however participants continued with the same mind-wandering instructions for the full 20 minutes. The first ten minutes of the meditation time block spent in focused attention on the breath was utilized to help prepare the subject to relax and deepen into their chosen meditation practice, and breath-focused awareness is a component preparatory stage of practice for practitioners of all three traditions assayed. By assessing the brain activity during this state we aimed to assess whether breath-focused meditation practice might demonstrate differences between practitioners of different traditions and controls as well as whether it might demonstrate more similarity between groups than specific meditation practices of different traditions. The order of tasks performed was counterbalanced. For consistency, the duration of the instructed mind-wandering block (IMW) was equal to the duration of the meditation block (including breath awareness).

The EEG data were processed using the EEGLAB open source software version 14 [11] running on MATLAB R2019a [1] under a Linux/Debian operating system. EEG data were first referenced to the channel average and then down-sampled from 1024 Hz to 512 Hz. A high-pass filter at 0.5 Hz was applied to remove signals related to head and body movements. We automatically removed portions of the signal and channels presenting non-stereotyped artifacts using the EEGLAB plugin `clean_rawdata` version 0.34, which uses Artifact Subspace Reconstruction [10]. The processed data were then checked visually for any potential remaining artifact. These recordings were also used in Vivot et al., 2020 [35] and Brabozscz et al., 2017 [3].

### 2.2. Meditation Techniques

#### 2.2.1. The Himalayan Yoga Tradition

The Himalayan Yoga Tradition (HT) is an ancient yoga tradition which focuses on the integration and purification of thoughts, emotions, mindfulness (including mindful breath awareness), mantras, postures, and body energies (chakras) [3]. This tradition is representative of those which incorporate mantras, as the primary meditation practice employed is the mental repetition of the mantra concurrently with breath awareness. Given the maintenance of focus on both mantra and breath, this practice represents a version of focused attention style meditation and we expected this specific meditation state to be relatively similar to breath awareness, meaning a short distance between the two meditations on the state space.

#### 2.2.2. Isha Yoga

Isha Yoga is currently maintained by the Isha Foundation, founded in 1992, and is composed of specific postures, breathing techniques, and sitting meditation [3]. Practitioners in the Isha Yoga tradition do many practices involving mantras, visualizations, and breath and body awareness. More advanced practitioners also learn the particular sitting meditation the participants in this study utilized: “Shoonya” meditation. Shoonya is translated as “emptiness,” “non-doing,” or “no-thingness.” Shoonya meditation, here abbreviated SNY, is engaged by Isha Yoga practitioners after preparation with various mantra and breath practices and involves the process of suspending or letting go of the operations of thinking. Due to this intentional separation from any particular thoughts within awareness it was hypothesized that the mental state during meditation might be most different from the mental state of breath awareness in this group. While Isha Yoga practitioners engage in a range of focused attention practices, the specific Shoonya practice assayed here is somewhat representative of the open-monitoring style of meditation, and it was predicted that the Shoonya meditation state would demonstrate large distances from the breath awareness state.

#### 2.2.3. Vipassana

Vipassana meditation, also known as insight meditation, is a form of meditation belonging to the Theravada tradition of Buddhism, reintroduced to the practice in the 18th century in Burma. It has since become popular in America and the basis for the development of western “mindfulness meditation” interventions. The Vipassana participants in this study practiced the form of Vipassana meditation taught by S.N. Goenka [19], which involves systemically scanning the sensations of the body after an initial period of breath awareness to calm and focus the mind. Vipassana practitioners in this tradition keep their attention moving throughout the practice, observing closely, objectively and with equanimity, the sensations he/she experiences [3]. It is of note that while some forms of Vipassana practice place a great deal of emphasis on the open-monitoring aspect of practice, the Goenka school of Vipassana is a focused attention form of Vipassana practice. Because Vipassana meditation is practiced with a strong focus on mindfulness of somatic sensations, it was hypothesized that the Vipassana meditation state would occupy nearby areas as the breath awareness state.

### 2.3. Dynamic Functional Connectivity

After the EEG was preprocessed to be artifact-free and appropriately filtered as detailed above, we created a dynamic functional connectome by calculating its functional connectivity via phase lag. This study utilizes weighted phase lag index (WPLI) [56], a measure of phase synchrony which circumvents the problems of volume-conduction and noise of the phase lag index [50], to construct the functional connectivity across time. The WPLI uses the imaginary component of the cross-spectrum to determine the phase leads and lags of two signals, as well as the magnitude of the imaginary component to weight phase differences. It is calculated as follows:

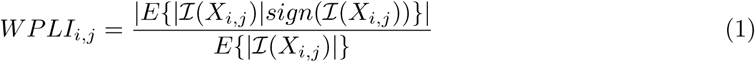

Where *E*{·} is the expected value, *X*_*i,j*_ is the cross-spectrum of channels *i* and *j, ℐ*(*·*) is the imaginary component, and *sign*(*·*) is the sign function [56]. In computing the WPLI, the FieldTrip MATLAB toolbox was utilized [39]. Each discrete frequency was calculated independently from 1 *–* 50 Hz using equation 1 implemented with the FieldTrip function `ft_connectivityanalysis` [51].

To create a representation of the dynamic functional connectivity, the continuous resting state EEG was organized temporally by a sliding window of length one second, which itself was divided into ten 0.1 sec epochs.

The window shifted 0.5 seconds forward each iteration, having thus a 50% overlap with the previous window. For each one second epoch, a 3D connectome, *C*^*s*^(*t*), was calculated using Definition 1.

**Definition 1**. Let *C*^*s*^(*t*) be the *Channel × Channel × Frequency* static functional connectivity matrix at time *t* for subject *s*, labeled a connectome.

Each task (e.g. meditation or IMW) is a set of *{t*_1_, *t*_2_, …, *t*_*end*_*}* from a single recording. For *t*_*i*_, *C*^*s*^(*t*_*i*_) is the static connectome formed using equation 1 at that time point with frequencies 1 *–* 50 Hz. To form the state space, the union of the tasks is taken to create a single set of time points. For simplicity, the task set is taken as a 4th dimension of the matrix formed in Definition 1, forming the dynamic functional connectivity (dFC) of that individual, *s*, with dimensions *Channel × Channel × Frequency × Time*.

### 2.4. Manifold Learning

This study uses a modified version of a manifold learning procedure called Thought Chart introduced in Xing et al, 2016 [61] and expanded here. From the dFC formed above, a dissimilarity matrix *D* was constructed by taking the Frobenius norm of the difference of each pair of connectomes, (*C*^*s*^(*t*_*i*_), *C*^*s*^(*t*_*j*_)), for all time points such that *D* has dimensions *t*_*max*_ *× t*_*max*_, as shown in equation 2.

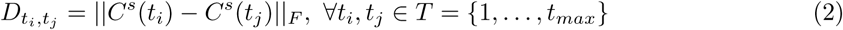

Each element of *D* gives a magnitude of the difference between the static connectomes, *C*^*s*^, at different time points in a matrix. The matrix *D* is also a fully connected graph in which each node is some *C*^*s*^(*t*) and its dissimilarity to each other connectome is encoded in its edges, to which manifold learning can be applied.

As *D* is fully connected, there exists connections which may not actually exist between connectomes. To address this, the *k* -nearest neighborhood (KNN) filtering algorithm is applied to *D*, with *k* defined as 5% of the total number of time points, or the minimum *k* such that *D* remains a connected graph. From this filtered matrix of local neighborhoods, we apply Dijkstra’s algorithm to find the geodesic distances between all *C*^*s*^(*t*), *t ∈ T* [54], as shown in figure 1. The resulting graph is a high-dimensional metric space which preserves distances between connectomics states, defined in Definition 2.

**Figure 1:**
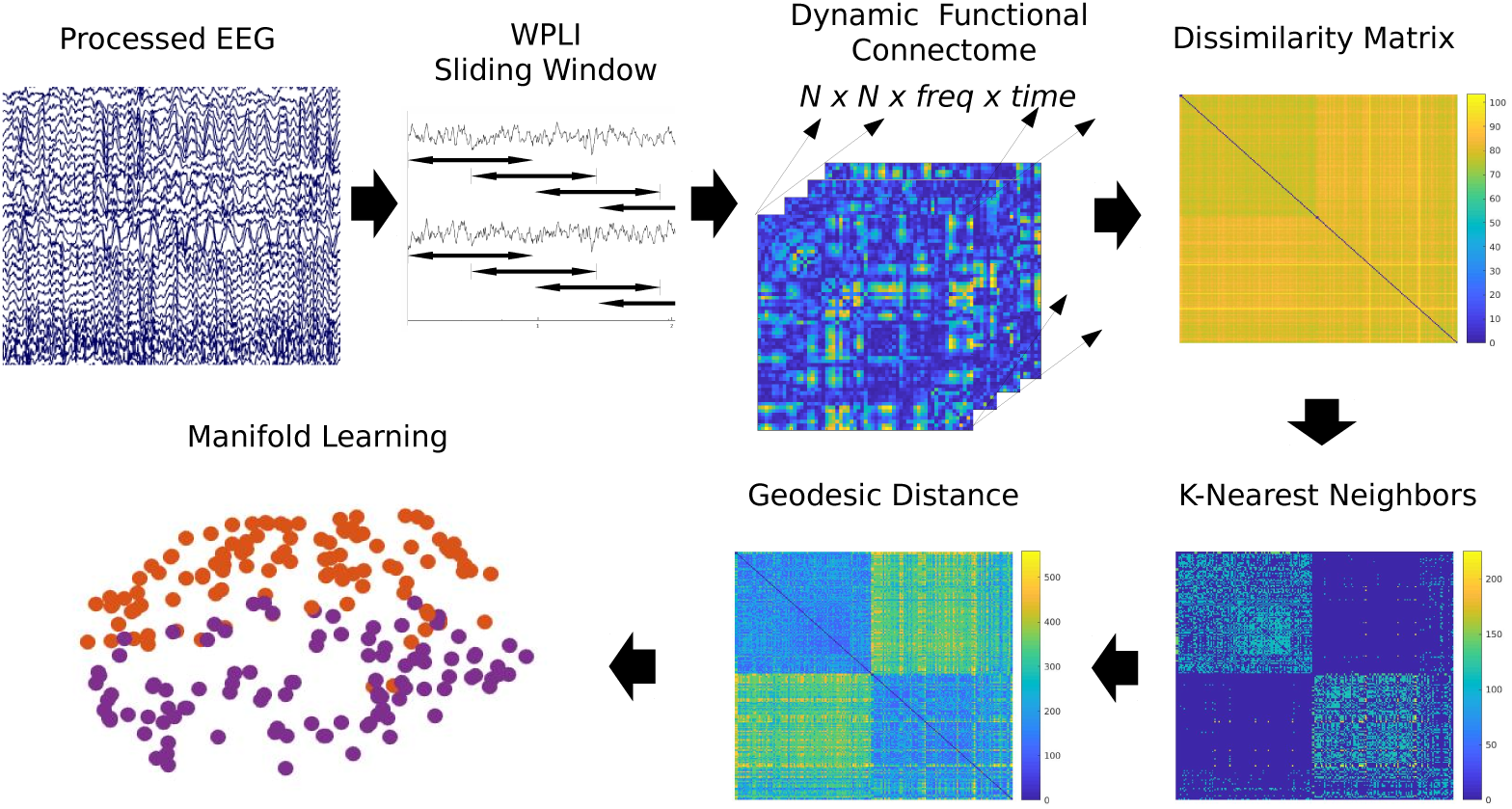
Overview of the Thought Chart method. The preprocessed EEG is passed through a weighted phase lag index algorithm to calculate a 3D (Chan, Chan, Freq) connectome at each time point (4D) during a given task, this figure shows two tasks. For each time point, the dissimilarity of every pair of 3D connectomes is determined and occupies one element of a dissimilarity matrix. This matrix then disconnects all but the closest 5% of neighbors, and the Dijkstra algorithm is used to find the true distance between each connectome on a manifold. That manifold can be visualized in a number of ways and comprises the Thought Chart of that individual.

**Definition 2**. Let the metric space 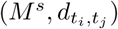, where *M*^*s*^ = *{C*^*s*^(1) … *C*^*s*^(*t*) … *C*^*s*^(*t*_*max*_)*}* and 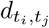 is the geodesic distance between *C*^*s*^(*t*_*i*_) and *C*^*s*^(*t*_*j*_), be labeled the Thought Chart.

This method is very similar to the algorithm for Isomap, a method of dimensionality reduction, which is used for visualization in this study as well [54]. This algorithm is illustrated in figure 1, and results in the Thought Chart for that individual, or the state space that encodes as distances, the similarity of connectomic states.

### 2.5. Hausdorff Distance

The Hausdorff distance is a metric used here as a measure of the similarity between two clusters of time points within a metric space [18]. As the connectomes at each time point become more similar to each other, they are closer together in the state space. Thus, when comparing two sets of *{C*^*s*^(*t*)*}*, as the similarity between the two tasks increases, the Hausdorff distance between them decreases. To account for potential scaling difference between subjects, the Hausdorff distances were normalized to the mean of all geodesic distances within that subject. The Hausdorff distance between subsets *A* and *B* in a metric space *M* is defined formally in equation 3.

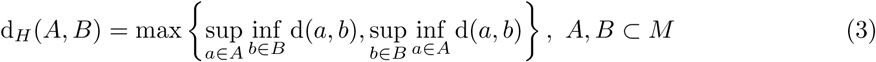

where sup is the supremum and inf is the infimum, which is then normalized by the mean *d* in *M*. Here, subsets *A* and *B* would each be a set of *{C*^*s*^(*t*)*}* during a specific task performed by the same individual.

### 2.6. Analysis and Statistics

The Thought Chart method was performed using MATLAB, R2019a [1], and the graph metrics were computed using functions found in the Brain Connectivity Toolbox [42]. The graph metrics were calculated independently for each task group, namely the meditative tasks (breath awareness and meditation) and IMW task. For the characteristic path length (CPL), global efficiency, betweenness centrality, and normalized trajectory length (NTL), a distance matrix based on dissimilarity between connectomes was used, normalized by its global mean distance; for the node strengths, assortativity, and clustering coefficient a connectivity matrix was used. The connectivity matrix was computed by subtracting each element in the distance matrix from the max distance, and then normalizing to be between 0 and 1, with the diagonal set to 0. This connectivity matrix was then thresholded proportionally, keeping 25% of edges.

All statistics were computed in R version 3.6.3 [53] and using packages `car` [14], `dplyr` [58], `tidyr` [59], `afex` [47], `emmeans` [28], `ggplot2` [57], and `R.matlab` [2]. Paired t-tests were used to determine differences between graph metrics in tasks. Two-Way ANOVAs were used to determine differences between groups and tasks, except where noted otherwise. Follow-up tests were performed using linear contrasts, with FDR corrections. Brown-Forsythe tests were used to test homogeneity of variance.

## 3. Results

### 3.1. Characteristics of the Thought Chart

The participants did vary in their demographics, presented in table 1, in both age (*F* (3, 67) = 3.21, *p* = 0.03) and years of meditation experience (*F* (2, 50) = 14.17, *p* < 0.0001. When applicable, these two variables are controlled for as covariates, in addition to sex.

**Table 1:** Demographic information about the participants, in the form of means and (standard deviations).

|  | Control | Himalayan | Isha | Vipassana |
| --- | --- | --- | --- | --- |
| Age | 44.4<br>(10.0) | 51.1<br>(13.9) | 39.1<br>(10.5) | 49.2<br>(18.2) |
| Years of Practice | 0 | 24.4<br>(13.9) | 6.9<br>(4.1) | 16.2<br>(11.6) |
| Sex (% female) | 38.8 (7/18) | 22.2 (4/18) | 11.1 (2/18) | 22.2 (4/18) |

Each individual whose EEG recording is preprocessed and analyzed using the Thought Chart method, illustrated in figure 1, produces their own unique state space to be analyzed further. Each participant’s Thought Chart for these tasks had a mean of 2434.4 *±* 62.8 time points, with 608.6 *±* 15.7 time points per task (each one second long). *k* for *k* -nearest neighbors was minimized for each person to keep their state space fully connected and for each participant was 222.2 *±* 149.6 nearest neighbors (or 9.1% of the mean time points).

### 3.2. Task separation occurs following Thought Chart analysis

After the manifold learning, detailed above, the resulting state space was visualized using Isomap as it adequately preserves geodesic distances. One such representative individual’s Thought Chart, visualized with Isomap, is shown in figure 2, however all analyses were done using the high dimensional data. For each task, the *C*^*s*^(*t*^*∗*^) where *t*^*∗*^ is the time point with the highest betweennesscentrality is shown with the frequencies averaged in order to illustrate what this connectome may look like in the standard *Channel×Channel* connectivity map generated from WPLI. The two IMW tasks (labeled as ‘IMW 1’ and ‘IMW 2’ and referring to either the first 10 minutes of IMW or the second 10 minutes of IMW respectively) are largely co-mingled together, as are the two meditation tasks (’Breathing Awareness’ and ‘Meditation’). We then measure the distance between the point clouds representing each task using Hausdorff distance to get a sense of how much separation there is for each person.

**Figure 2:**
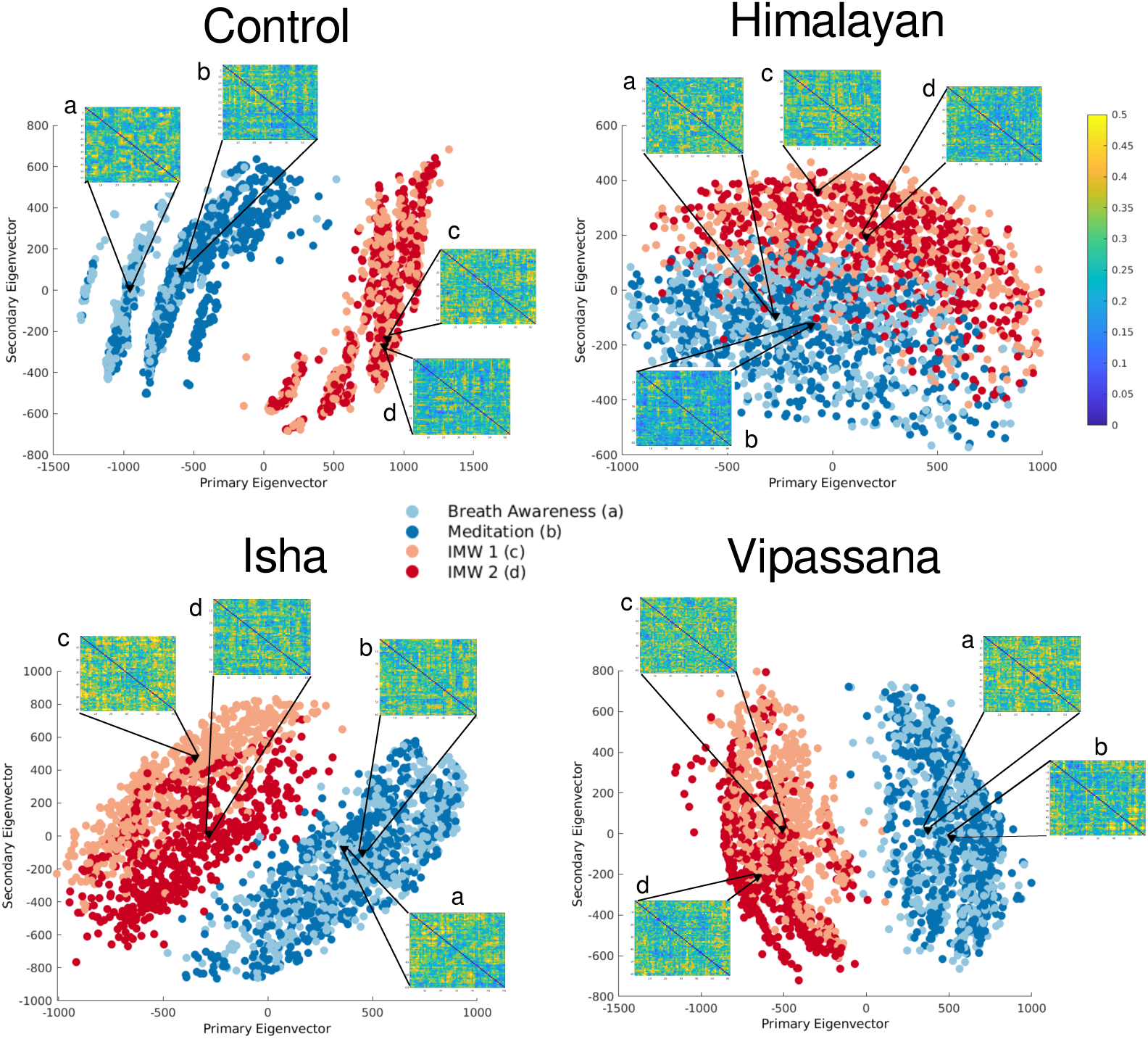
The Thought Chart of four representatives from each groups (Control, Himalayan, Isha, and Vipassana), visualized using Isomap, demonstrating the task separation in the state space. For each representative, the central-most time point *C*^*s*^(*t*) for each task is visualized as an EEG connectivity matrix, with frequency dimension averaged. The four tasks are (a) Breath Awareness, (b) Meditation, (c) Instructed Mind-Wandering 1, and (d) Instructed Mind Wandering 2. All EEG connectivity matrices share the color bar on the right from 0 to 0.5, which is the strength of the relationship as measured by WPLI.

For more quantitative analyses of the task separation see figure 3. As each task was a separate EEG recording, acquired in succession while the participant stayed engaged in their task, we aimed to determine whether this was artifact or represented real differences in the mental landscape of the tasks themselves.

**Figure 3:**
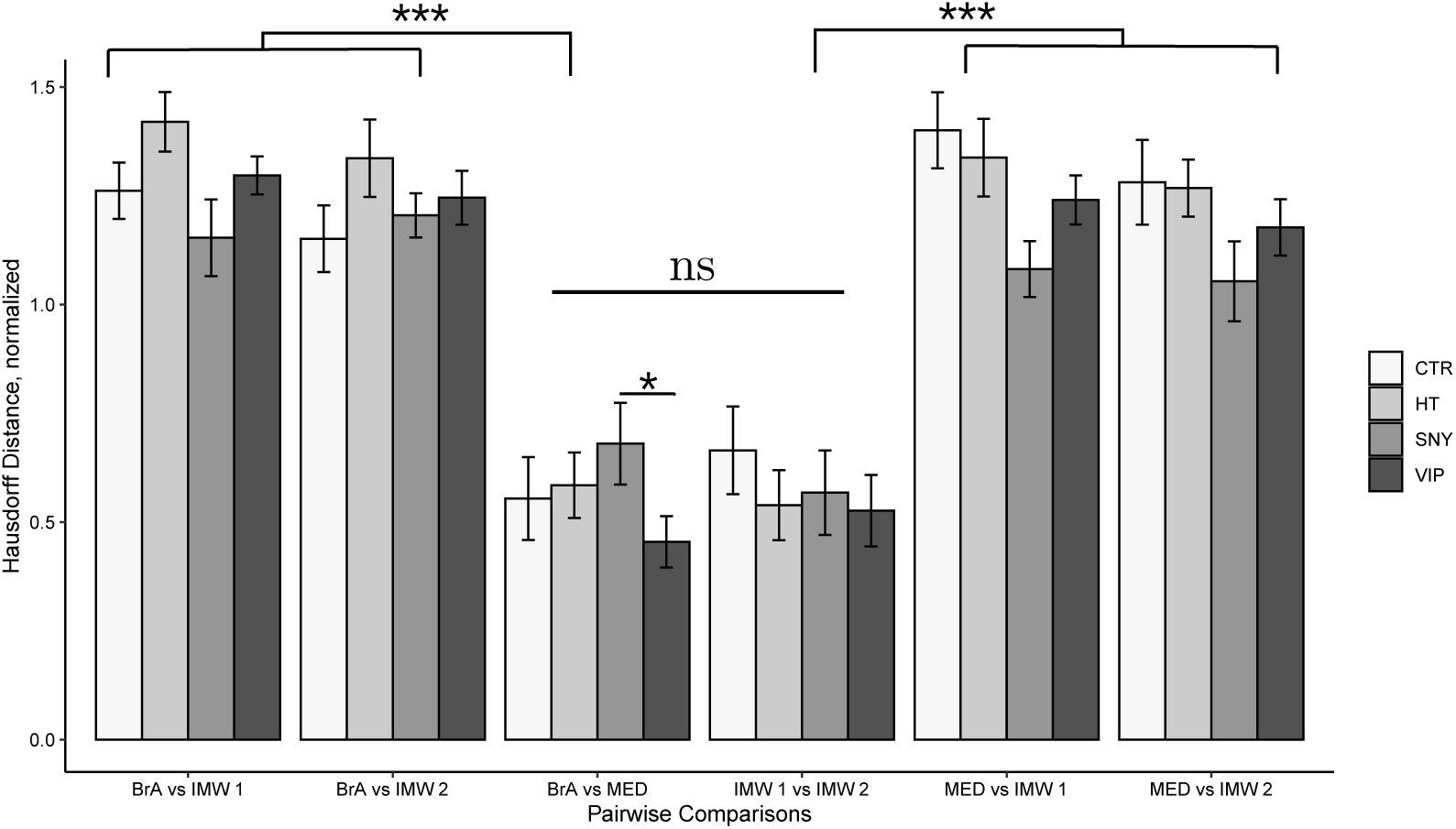
Bar plots of the Hausdorff distances of each pair of tasks. Each bar from left to right corresponds to controls, Himalayan Tradition, Isha Yoga, and Vipassana. The y-axis is the Hausdorff distance between the two tasks being compared in the x-axis, normalized to the mean geodesic distance. The tasks along the x-axis are breath awareness (BrA), meditation (MED), and instructed mind-wandering (IMW 1 & IMW 2). The two IMW tasks were significantly closer together than the unrelated pairs of tasks as were the two meditative tasks. *ns*: *p* > 0.05, \**p* < 0.05, \*\*\**p* < 0.0001.

### 3.3. Cluster separation is due to task differences

To evaluate whether the separation shown in 2 is due to variations due to the time the recording took place or differences in the dynamics of the task, we compared the state space distance between each pair of tasks. If the separation in the state space is due to the differences in the tasks, one would expect the two recordings of the same task to be sufficiently similar to each other in a given individual, while the distance between different tasks would be significantly greater. Thought Chart was applied to four recorded tasks for each individual and the pairwise proximity of tasks determined using Hausdorff Distance, described above [27]. The results of these comparisons are shown in figure In all groups of meditation, the similar meditation tasks (breath awareness vs. specific meditation) and the similar IMW tasks (IMW 1 and IMW 2) were very close to each other, with a mean Hausdorff distance near half of the average distance (*M* = 0.574 ± 0.032 for two IMW tasks and *M* = 0.568 ± 0.031 for the two meditation tasks). The distances between dissimilar tasks were above the average geodesic distance for all groups as demonstrated in figure 3.

A two-way ANOVA showed that the pairwise distances were significantly different by which two tasks were being compared, but not by meditation group, *F* (3.2, 217.7) = 78.98, *p* < 0.0001, with Greenhouse-Geisser corrections. A Box’s M-test for homogeneity of covariance was not significant, *p* = 0.31. Bonferroni-corrected follow-up paired t-tests showed the inter-task (IMW vs. meditation) distances were significantly greater than the intra-task distances, all with *p* < 0.0001, and the difference of mean distance of the meditation tasks and the IMW tasks were not significantly different. The comparison of breath awareness vs. meditation was hypothesized to show that in the Vipassana group the two tasks would be closer in the state space than in other groups, particularly the Shoonya meditation, due to the importance of breath awareness during Vipassana and de-emphasis in Shoonya. Indeed, the Vipassana group had a significantly shorter Hausdorff distance between these two meditative tasks than the Isha (SNY) group, *t*(68) = 2.65, *p* = 0.048, though neither were significantly different from the control and an ANOVA showed that the difference between groups over this pair of tasks is merely near significant when all four groups are considered, *F* (3, 68) = 2.39, *p* = 0.08. There was no significant difference between the groups in the meditation vs. IMW (IMW 1 and IMW 2) comparisons, *F* (3, 68) = 1.89, *p* = 0.14.

### 3.4. Task differences do not rely on a specific frequency band

To determine the extent to which the cluster of mental states (*{C*^*s*^(*t*)*}*) incorporating a certain task in the Thought Chart of an individual is driven by a particular frequency band, a pair of post-hoc complementary analyses were conducted using a subset of frequencies during the manifold learning step. Specifically, for each individual, in the first analysis of each of the frequency bands, delta (1 *–* 3 Hz), theta (4 *–* 7 Hz), alpha (8 *–* 14 Hz), beta (15 *–* 31 Hz), and gamma (32 *–* 50), were isolated during the manifold learning step (excluding all other frequencies) while in the second analysis all frequency bands except one were excluded for manifold learning. Then, the Hausdorff distance between each pair of tasks were calculated. For simplicity, we computed only results for the control group for this analysis. As indicated in figure 4, the properties of the task separation remains in both the isolation and the exclusion frequency analysis. A two-way repeated measure ANOVA showed that when considering the pairwise task differences, only the comparisons themselves were significant, *F* (5, 10) = 11.75, *p* < 0.001, whereas the frequency bands and whether the band was isolated or excluded, and any interaction between these variables were not significant, all *p* > 0.05. Follow-up contrasts results showed that the same pattern as figure 3 holds for all frequency bands, excluded or isolated. The intra-task Hausdorff distances (IMW and breath awareness/meditation) were significantly lower than the inter-task Hausdorff distances, *t*(10) = -7.18, *p* = 0.0001, while the two intra-task Hausdorff distances were similar to each other, *t*(10) = 1.01, *p* = 0.3.

**Figure 4:**
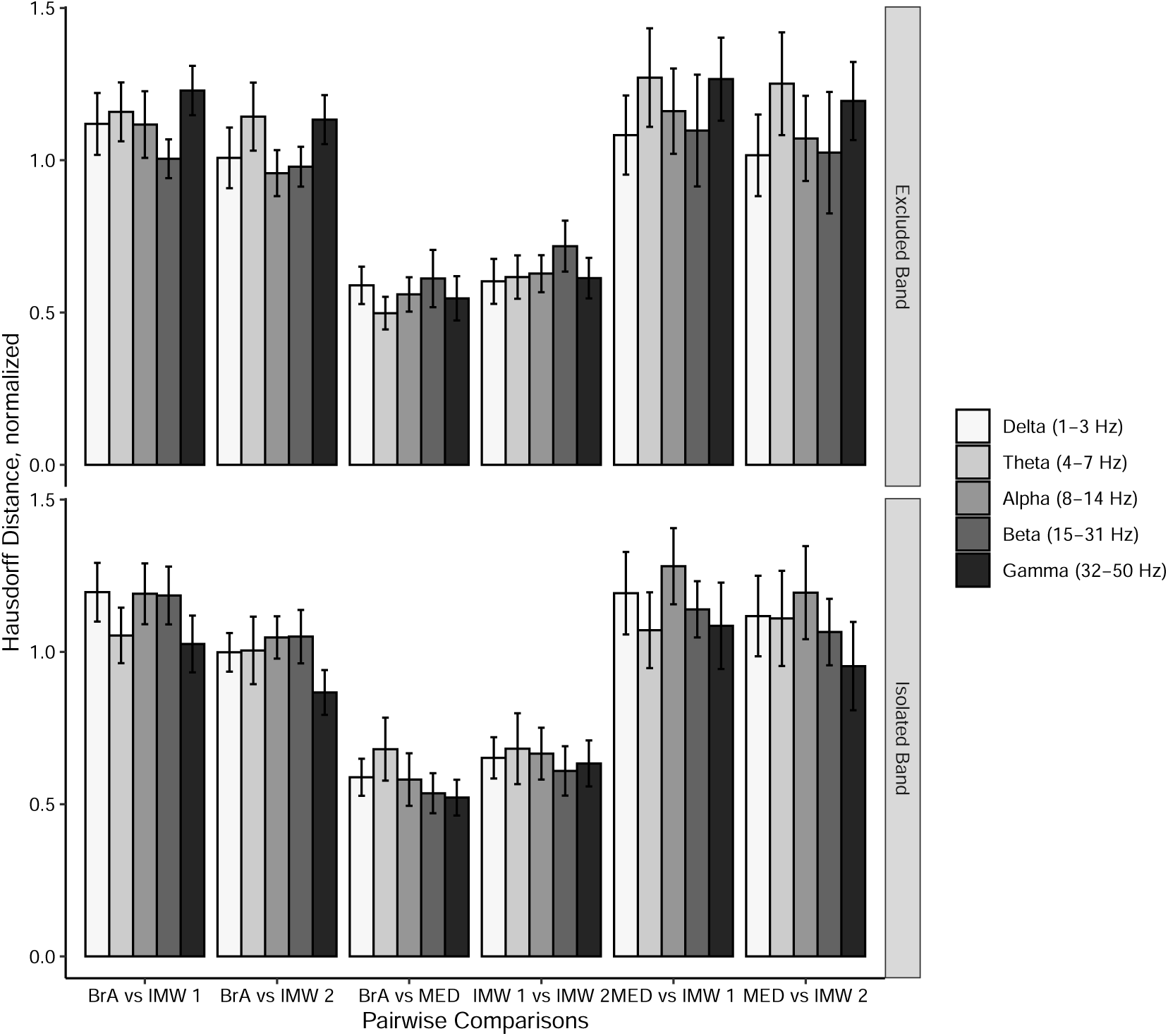
The mean Hausdorff distance between each task, error bars showing standard error. The upper figure indicates the normalized Hausdorff distance between tasks where each band is excluded, and the lower figure shows the same with frequency bands isolated. In both, inter-task distances were significantly different while intra-task were not. BrA = Breath Awareness, MED = Meditation, IMW = Instructed Mind Wandering.

### 3.5. Graph Metrics of Thought Chart

To further describe the state space, table 2 shows some of the global graph metrics of the final resulting Thought Charts, separated into the two classes, meditation and IMW, tasks. CPL, efficiency, centrality, and NTL were computed using a normalized distance matrix, while node strength, assortativity, and clustering coefficient were computed using a normalized thresholded connectivity matrix. Paired t-tests did not detect differences between meditations and IMW in any metric. Using two-way ANOVAs to find differences between groups and tasks showed no significantly deviating groups in the graph metrics calculated here.

**Table 2:**
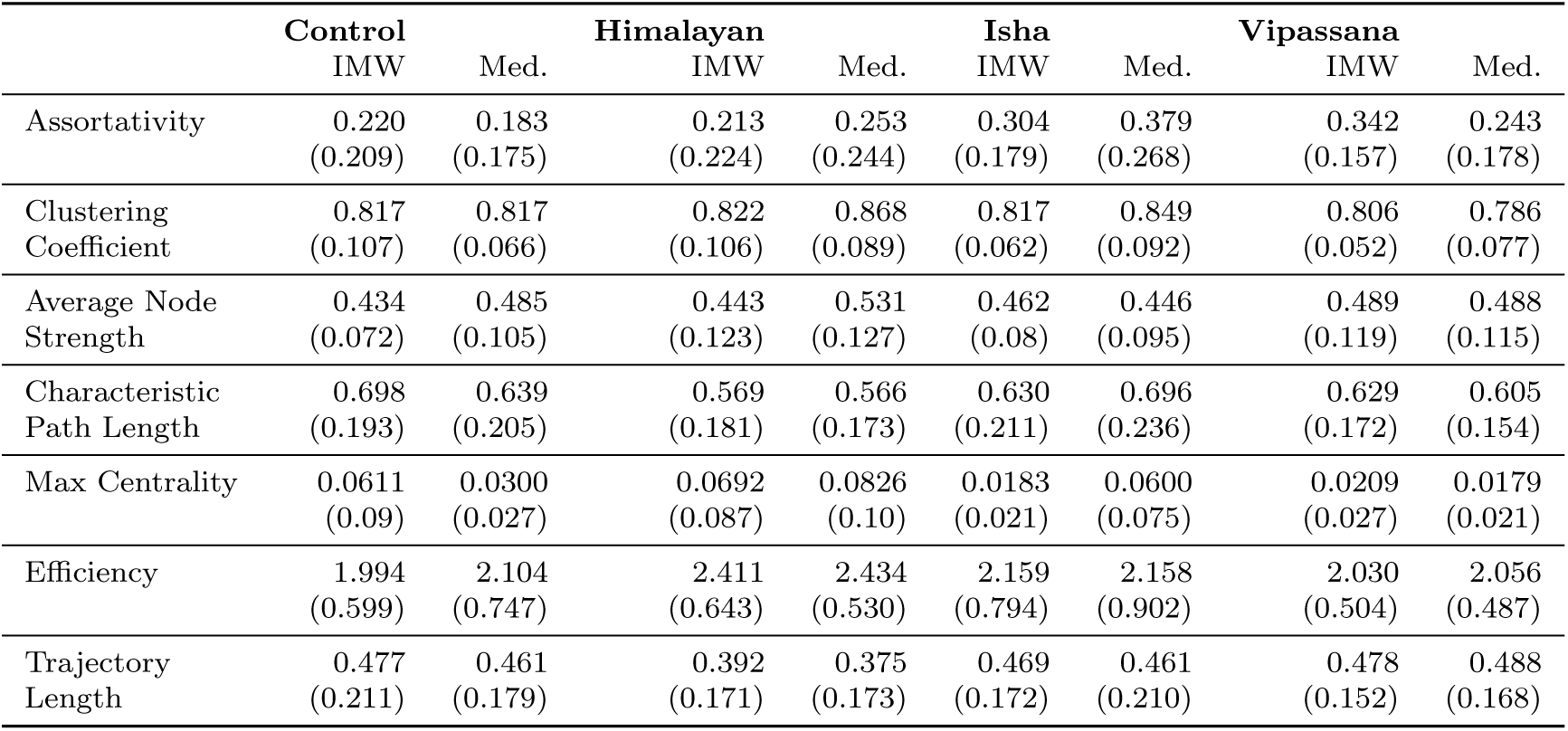
Some global graph metrics of the Thought Chart state space, in means and (standard deviation), for each task group and meditation group. Assortativity, Mean Node Strength, and Clustering Coefficient were computed on a binarized thresholded connection matrix. CPL, Centrality, Efficiency, and Trajectory were computed used a weighted distance matrix.

|  | **Control** |  | **Himalayan** |  | **Isha** |  | **Vipassana** |  |
| --- | --- | --- | --- | --- | --- | --- | --- | --- |
|  | IMW | Med. | IMW | Med. | IMW | Med. | IMW | Med. |
| Assortativity | 0.220<br>(0.209) | 0.183<br>(0.175) | 0.213<br>(0.224) | 0.253<br>(0.244) | 0.304<br>(0.179) | 0.379<br>(0.268) | 0.342<br>(0.157) | 0.243<br>(0.178) |
| Clustering Coefficient | 0.817<br>(0.107) | 0.817<br>(0.066) | 0.822<br>(0.106) | 0.868<br>(0.089) | 0.817<br>(0.062) | 0.849<br>(0.092) | 0.806<br>(0.052) | 0.786<br>(0.077) |
| Average Node Strength | 0.434<br>(0.072) | 0.485<br>(0.105) | 0.443<br>(0.123) | 0.531<br>(0.127) | 0.462<br>(0.08) | 0.446<br>(0.095) | 0.489<br>(0.119) | 0.488<br>(0.115) |
| Characteristic Path Length | 0.698<br>(0.193) | 0.639<br>(0.205) | 0.569<br>(0.181) | 0.566<br>(0.173) | 0.630<br>(0.211) | 0.696<br>(0.236) | 0.629<br>(0.172) | 0.605<br>(0.154) |
| Max Centrality | 0.0611<br>(0.09) | 0.0300<br>(0.027) | 0.0692<br>(0.087) | 0.0826<br>(0.10) | 0.0183<br>(0.021) | 0.0600<br>(0.075) | 0.0209<br>(0.027) | 0.0179<br>(0.021) |
| Efficiency | 1.994<br>(0.599) | 2.104<br>(0.747) | 2.411<br>(0.643) | 2.434<br>(0.530) | 2.159<br>(0.794) | 2.158<br>(0.902) | 2.030<br>(0.504) | 2.056<br>(0.487) |
| Trajectory Length | 0.477<br>(0.211) | 0.461<br>(0.179) | 0.392<br>(0.171) | 0.375<br>(0.173) | 0.469<br>(0.172) | 0.461<br>(0.210) | 0.478<br>(0.152) | 0.488<br>(0.168) |

One notable metric of the Thought Chart, is that when thresholding to the strongest 25% of connections and binarizing, there is a high clustering coefficient (between 0.79*–*0.87 on average), and assortativity was moderate in most groups (0.22 *–* 0.379 on average). Of the distance matrix metrics, the CPL was near half of the mean edge length of the full state space (between 56.6% *–* 69.8% of the mean edge length).

## 4. Discussion

In this study, we have investigated three groups of meditation practitioners using the Thought Chart method and found that in each group, as well as meditation-naïve controls, each meditative brain state occupied a different area of the state space than the mind-wandering tasks, see figure 3. This separation has also been shown to be independent of which frequency bands were included in the analysis (figure 4). We have also described some of the network characteristics of these Thought Charts for reference, see table 2. The high clustering coefficient and low characteristic path lengths indicate that this state space has characteristics of a small-world network, which was expected. States tend to cluster together when they are similar, with lower than random connections between clusters. The Thought Charts also have moderate positive assortativity, meaning that states with similar characteristics such as degree are more likely to be connected than random chance [13].

The work presented here characterizes a high-dimensional state space in which functional connectomes are neighbors to each other if they are similar, defined by the dFC of a single individual, called the Thought Chart. From this, we conclude that Thought Chart is capturing both the similarities and differences of the mental state in these meditation states and non-meditative tasks, and thus is a potential framework for analyzing them. Such a space has been described previously, but not in the context of healthy individuals [61, 24] and not on the individual level. We find that the shape of the Thought Chart for each individual varies, likely due to the significant between-subject variability in connectivity [41, 34] as well as potential contributions related to unique personal styles of meditation practice even within the similar meditation traditions.

We also showed that the separation in the state space of breath awareness task and the IMW task was not reliant on meditation experience, as the controls had similar separation as the experienced meditators. This was unexpected and could indicate a particularly low barrier to meditation. We speculate that one reason may be due to the stark differences in the tasks that the participants were asked to complete, one (IMW) concerned largely with non-focused reminiscence and the other (breath awareness) requiring strong focus and awareness of the present, even among the experienced practitioners. An alternative reason may be that the meditation-naïve participants successfully engaged in the meditation task during their breath awareness task, or at least were able to achieve a mental state sufficiently dissimilar to IMW to be indistinguishable from the experienced meditators from the three traditions studied. If the latter is the case, then we believe these results show that everyone can learn to meditate, from our healthy sample in this study.

The Thought Chart shows promise as a method to easily and affordably map the relative locations of mental states in individuals’ mental state space using EEG, and this utility does not seem to be much different for meditators vs. non-meditators. Alpha and gamma frequencies may play an important role in meditation, as shown in Braboszcz et al. [3] analysis of this same dataset, wherein all meditation groups’ practitioners demonstrated a state-trait increased high gamma power in the parieto-occipital region as well as specifically increased trait alpha power in the Vipassana meditator group. Nonetheless, as indicated in figure 4, Thought Chart discriminability does not rely on a particular frequency band, a finding that was true across all four groups. Initial analysis of this dataset utilizing the ThoughtChart approach found that due to great between-individual differences in dFC and resultant ThoughtChart metrics, group-level analyses did not yield much in the way of significant group differences. Interestingly, within-individual differences in brain state were much more sensitively identified in this dataset, even clearly distinguishing the thinking/mind-wandering states from the meditation states in meditation-naïve participants. We do not yet know whether Thought Chart can be performed using fMRI BOLD recordings instead of EEG, however we suspect that the temporal resolution of EEG is of great importance to the accurate mapping of the thought state space. Future work on this topic will look at its use in more tasks as well as using different techniques to construct the connectome from the EEG time series such as mutual information or Granger causality as these have also been shown to have a high degree of individual identifying information [45]. Confirming that these methods produce similar Thought Charts would provide further supporting evidence of the results herein.

Our group has previously demonstrated the utility of some universally applicable state space on the group level [61, 62]. Xing et al. were able to describe this space using the data from groups of individuals performing the Emotion Regulation Task (ERT), with different trajectories in the group’s thought space in individuals with social anxiety, indicating the similarity of the location of these mental states across subjects. An important and necessary improvement on those methods were to expand its functionality to the individual level, due to the known inherent differences between each person’s functional connectivity, even among healthy people [45]. As this metric space was constructed using these relative differences as its basis, it makes sense to employ the Thought Chart method on the group level to explore group-based differences, but applications such as meditation may be too noisy for group-level analyses and therefore this within-individual Thought Chart can now be utilized. It is also often clinically useful to analyze individuals instead of groups, as the rise of personalized medicine, particularly in psychiatry, indicates [40], and this new form of Thought Chart may be useful in those domains as well.

One potential reason for these task separations in the state space is differences in the ratio of time spent in different microstates [64]. Though it could be a major driver of the separation, we do not think it to be the only source of these state differences, as in that case, we would expect to see more overlap, as the same microstates are being used for different tasks. Further research would be needed to demonstrate to what extent these results depend on microstate differences captured by EEG [37].

Though we did not find any significant trait differences between the four groups in this study, the use of a diverse group of meditators allowed us to demonstrate the broad use of these methods in healthy people with known differences in functional connectivity [20, 52]. We were also able to show, as per our hypothesis, that Vipassana meditation was closer to breath awareness in the Thought Chart analysis than Shoonya meditation was, possibly reflecting that there is a greater similarity between breath-focused meditation and this specific style of body-awareness focused Vipassana practice compared to the “emptiness/no-thingness” practice of Shoonya. It is interesting to note that every tradition of meditation tested demonstrated both a similarity to breath awareness on a subject-by-subject level of analysis, and a dissimilarity to autobiographical mind-wandering. The within-individual Thought Chart is not designed to find differences between groups, for this one should consider the original Thought Chart [61] method. Here, the within-individual formulation of Thought Chart can acutely sense and provide a method of visualization of the difference between tasks or states within individuals. We believe that this tool can be utilized to explore tasks which would not trigger strong or acute responses, but subtler temporal patterns. In this paper we have shown that meditation is distinctly different from mind-wandering among both experienced and naïve meditators, as determined by their different position in the state space. The objective quantifiable difference is measured by the Hausdorff distance between those positions, which is supported in the literature when meditation is compared with resting-state [6, 52]. We have also shown that these differences utilize the entire connectomic frequency space, from 1 *–* 50 Hz, and are not dependent on a particular frequency band.

Extensive previous evidence demonstrating different brain functional connectivities distinguish long term practitioners from controls and previous studies demonstrate that the meditator groups comprising the current dataset demonstrate enhanced high gamma power [3] and increased sample entropy [35] relative to the meditation-naïve participants. These studies found some specific differences between Vipassana meditators in particular and the other groups, both enhanced alpha power and the greatest degree of entropy [35]. Thus, future analysis of this data utilizing the original Thought Chart method might be usefully applied to help determine whether other differences can be identified between groups given that the long hours of meditation within the three different traditions are likely to have had substantial differential effects on network connectivities.

In conclusion, individualized Thought Chart presents a compelling method for comparing the temporal dynamics of functional connectomes in a state space. Future work on Thought Chart may reveal it to also be a promising candidate for predictive or diagnostic biomarker for psychiatric disorders and treatment progression.

